# The cryptic step in biogeochemical Tellurium (Te) cycle: Indirect elementary Te oxidation mediated by manganese-oxidizing bacterium (MnOB)

**DOI:** 10.1101/2023.02.22.529621

**Authors:** Yuqing Liu, Huiqing Ma, Ang Li, Xianliang Yi, Yang Liu, Jingjing Zhan, Hao Zhou

**Affiliations:** Key Laboratory of Industrial Ecology and Environmental Engineering (Ministry of Education), School of Ocean Science and Technology, Panjin Campus, Dalian University of Technology, China; State Key Laboratory of Urban Water Resource and Environment, School of Environment, Harbin Institute of Technology, Harbin 150090, People’s Republic of China

**Keywords:** Manganese oxidation, Tellurite reduction, Biogeochemical cycle, Bacteria, Tellurium oxidation

## Abstract

Tellurium (Te) is a rare element in the chalcogen group, and its biogeochemical cycle has been investigated for decades. As the most soluble Te species, tellurite (Te(IV)) possess the highest toxicity to the organisms. Chemical or biological Te(IV) reduction to elemental tellurium (Te^0^) is generally considered as an effective detoxification route for Te(IV)-containing wastewater. Here, we reported a previously overlooked Te^0^ oxidation process mediated by manganese-oxidizing bacterium *Bacillus* sp. FF-1. This strain has both Mn(II)-oxidizing and Te(IV)-reducing activities, which could produce manganese oxides (BioMnOx) and Te^0^ (BioTe^0^) when incubating with Mn(II) and Te(IV), respectively. Te(IV) can co-precipitated with Mn(II) to form highly stable Te(IV)-Mn(II) compounds with low bioavailability. While when 5 mM Mn(II) was added after incubating 0.1 mM or 1 mM Te(IV) with strain FF-1 for 16 hours, the BioTe^0^ were certainly re-oxidized to Te(IV) by BioMnOx according to the results of X-ray photoelectron spectra (XPS) and Transmission electron microscope (TEM). The chemogenic and exogenous biogenic Te^0^ can also be oxidized by the BioMnOx, although with different rates. This study highlights a new transformation process of tellurium species mediated by manganese-oxidizing bacteria, revealing that the environmental fate and ecological risks of Te^0^ needed to be re-evaluated.

**Importance:** Biogeochemical cycle of Te mediated by bacteria mainly focus on the Tellurite reduction and methylation. In this study, the indirect tellurium (Te^0^) oxidation driven by manganese-oxidizing bacterium is firstly confirmed. As Te0 usually considered as a stable and safe products during Te(IV)-containing wastewater treatment, we suppose the ecological risks of Te^0^ needed to be re-evaluated due to the possible oxidation by manganese-oxidizing bacterium and its generated manganese oxides.

## Introduction

Tellurium (Te) is chalcogen metalloid belonging to group VIA with Oxygen (O), Sulfur (S), Selenium (Se) and Polonium (Po).^1^ Although the abundance of Te in earth crusts is low (~5 μg/kg), it has come into prominence due to the widely industrial application in solar panels, battery materials, petroleum refineries, and thermoelectric devices.^2^ During the production and usage of these materials, Te-containing species are inevitable released into the environment and caused ecological risks. In electroplating wastewater, the tellurium concentration can even reach 10 mg L^−1^.^3^ The most common Te species in environment including tellurate (Te(VI)), tellurite (Te(IV)), tellurium (Te^0^), and tellurides (Te(-II)). Te(IV) and Te(VI) are the dominant species under oxic conditions with high solubility. Compared with Te(VI), Te(IV) has higher solubility and more toxic to most of the microorganisms. The minimum inhibitory concentration (MIC) of Te(IV) to *Escherichia coli* was as low as 1 μg L^−1^.^4^ However, Te^0^ is generally insoluble and with low bioavailability, which results in low toxicity to organisms than Te oxyanions.^2^ Therefore, converting Te(IV) into Te^0^ is an efficient route to decrease the ecological risks of Te(IV) and recovery the scare Te resource.

Up to date, electrochemical, chemical, and biological methods have been employed to facilitate Te(IV) reduction in wastewater while only biological methods are preferred for field-level Te(IV) remediation due to the low cost.^5^ Various Te(IV) reducing bacteria have been identified and both enzymatic/nonenzymatic-driven Te(IV) reduction pathways were elucidated.^6–8^ In community level, Te(IV) removal by the bioabsorption and reduction capacities of aerobic/anaerobic activate sludge, granular sludge, and biofilm in membrane devices were also reported, with the generation of nanoscale Te^0^ (BioTe^0^) as the terminal products in most of the cases.^3, 9, 10^ The BioTe^0^ is thought to have high stability in environment, but several bacteria can further reduce BioTe^0^ into gaseous methyl tellurides species with much slow rate than Te(IV) reduction into Te^0^.^11, 12^ Compared with bacterial-driven Te^0^ reduction, whether Te^0^ can be oxidized by bacteria either directly or indirectly is rarely studied. In acidic condition (pH < 3.0), telluride (Te(-II)) in mine tailings can be oxidized to Te(IV) and Te(VI) in the presence of Fe(III) and bacteria *Acidithiobacillus ferrooxidans* through biogenic sulfuric acid, suggesting the possibility of Te^0^ oxidation in this condition.^13^ However, the Te^0^ produced in Te(IV)-containing wastewater usually in neutral condition, whether the Te^0^ oxidation could occur widely in the environment is still a mystery.

Manganese oxidizing bacterium (MnOB) can oxidize Mn(II) into manganese oxides through directly or indirectly processes.^14^ The biogenic manganese oxides (BioMnOx) are high reactive to reduced species, such as Cr(III), Co(II), Ce(III), and organic compounds due to the strong oxidation power of Mn(III)/Mn(IV).^15–18^ In addition, the lattice defect and large surface area also made the outstanding absorption capacity of BioMnOx to various metal ions.^14^ Nowadays, the synergy of MnOB and *in-situ* produced BioMnOx on organic pollutants and metal(loid)s oxidation were found. For Mn(II)-oxidizing fungus, the coupled Mn(II) and Cr(III) oxidation were found in three *Ascomycete* fungus. The effect of Cr(III) on Mn(II) oxidation was depend on the specific Mn(II) oxidation mechanism.^19^ In addition, simultaneously aerobic Se(IV)/Se(VI) reduction and Mn(II) oxidation occurred in two ascomycete fungus, and the volatile Se(-II) species could be quickly oxidized by mycogenic MnOx.^20^ However, whether Se° could be oxidized by BioMnOx was unconfirmed.

The objective of this study was to investigate the possibility of Te^0^ oxidation by MnOB and its derived BioMnOx. One MnOB with both Mn(II)-oxidation and Te(IV) reducing abilities was employed to check the *in-situ* Te^0^ oxidation by BioMnOx in different incubation conditions with Mn(II) and Te(IV). Furthermore, the oxidation of chemogenic and biogenic Te^0^ produced by other bacteria by strain FF-1 were also investigated. This study confirmed a cryptic Te^0^ oxidation step in the biogeochemical Te cycle, which was related to the BioMnOx produced by MnOB. Considering the widespread existence of MnOB in natural waterbody and wastewater treatment plants, this study provides a new insight of the environmental fate of Te^0^ and reveals the importance of re-evaluate the ecological risks of Te^0^.

## Materials and methods

### Chemicals and Bacteria strains

Leucoberbelin blue (LBB, 65%) was purchased from Sigma-Aldrich (USA), sodium tellurite was purchased from Aladdin Co. Ltd. (Shanghai, China). The other chemicals and reagents were purchased from Damao Co. Ltd. (Tianjin, China). All chemicals’ solutions were sterilized by filtration through a 0.22 μm filter membrane before use. The strains used in this study includes *Bacillus* sp. FF-1, *Bacillus cereus* CC-1 (CICC 24251) and *Aurantimonas* sp. HBX-1 (CGMCC 1.19345), which were isolated from surface seawater and marine sediments by our laboratory. Strain FF-1 and HBX-1 were previously isolated as Mn(II)-oxidizing bacteria (MnOB) and strain CC-1 as a Te(IV)-reducing bacteria.^21, 22^ The medium used for bacterial incubation was provided in supporting materials.

### Te(IV) and Mn(II) transformation experiments

For strain FF-1, it was incubated in 500 mL conical flasks containing 250 mL sterilized PYE medium with 1% (v/v) inoculum. The incubation was kept at 30°C, 150 rpm and without light irradiation. To evaluate the Te(IV) tolerance of strain FF-1, appropriate amount of Te(IV) (5 to 100 μM) was added into the medium after filtered sterilization. OD_660_ of the medium at specific time intervals was determined using an UV-vis spectrophotometer. The sustained Mn(II) oxidation ability of FF-1 has been confirmed in our previous study. The Mn(II) resistance of this strain could up to 7 mM and the optimal Mn(II) oxidation concentration was 5 mM.^14^ The initial Mn(II) concentration was set as 3 mM or 5 mM in this study, based on the consideration of growth rate of strain FF-1. The possible effects of the *in-situ* produced BioMnOx on the BioTe^0^ transformation were investigated under three conditions: (1) Mn0Te16h: 3 mM Mn(II) was added first with strain FF-1 while inoculating, then 1 mM Te(IV) added after 16 h; (2) Mn0Te0h: 1 mM Te(IV) and 3 mM Mn(II) added simultaneously with the bacteria; and (3) Te16Mn46h: 1 mM Te(IV) added 16 h after bacteria inoculation, then 5 mM Mn(II) added after 30 h. At specific time intervals, the medium was centrifugated (10000 rpm, 10min) to obtained the bacterial cells and Mn(II)/Te(IV) transformation products. The supernatant was filtered with 0.22 μm filter membrane and stored at 4°C for the residual total Te determination.

For comparison, chemical synthesized Te^0^ and BioTe^0^ produced by *Bacillus cereus* CC-1 were also prepared to compare the reactivity with BioMnOx produced by strain FF-1. The BioMnOx was obtained by incubating strain FF-1 with 5 mM Mn(II) for 6 days. Chemogenic Te^0^ was obtained by the reaction of 1 mM sodium tellurite with 5 mM cysteine for 20 min at magnetic stirrer (700 rpm).^23^ The products were centrifugated at 12,000 rpm and washed with ultrapure water for 3 times, and then dried at vacuum ovens (50 °C). The BioTe^0^ produced by strain CC-1 was done by adding 1 mM Te(IV) into pre-incubated bacterial cells for 24 h. After extra 5 days’ incubation, the products were collected by centrifugation, and used either directly (BioTe^0^ with active bacterial cells) or after sterilization at 121 °C for 20 min (inactive BioTe^0^). The sterilized BioTe^0^ was used to simulate the BioTe^0^ released into the environment after bacteria lysis. All the above experiments were conducted in triplicates and the results were presented as mean ± standard derivation (SD) value.

### Analytical Methods

Total Te concentration in supernatant were determined by ICP-MS. All samples in the above experiments were collected using sterile syringes and immediately filtered using 0.22 μm membrane filters, then diluted to ppm level with 2% (v/v) HNO_3_. The amount of BioMnOx at specific time intervals were determined using LBB method. The sample was mixed with LBB in a ratio of 1:5 and reacted for 5 min, centrifuging (11000 rpm, 5 min) and adding 180 μL of supernatant dropwise to a 96-well plate, measuring OD_620nm_ and calculating BioMnOx concentration using potassium permanganate as standard chemical.^24^ Gas chromatography (GC7890B, Agilent) equipment with sulfur chemiluminescence detector (SCD) was used to detect the possible gaseous methyl tellurides species in the headspace of cultures after 120 h incubation with a detection limit of 60 ppb (For dimethyl tellurides).^25^ For the annotation of the bacterial genome, MMseqs2 was used to identify the potential genes related to tellurite methylation using the thiopurine methyltransferase bTPMT from *Pseudomonas syringae* (Genbank No. AAC27664.1) as query sequences.^26^ The whole genome sequence of strain FF-1 has been deposited in the GenBank with an accession number of JAQQGG000000000. The version described in this paper is version JAQQGG010000000.

The morphological characteristics of the solid products were analyzed by scanning electron microscopy (SEM, Tecnai G2F30 STWIN, FEI, Inc., USA) and transmission electron microscopy (TEM, Nova Nano SEM 450, FEI, Inc., USA). For SEM, the bacteria cells and solid products were collected by centrifugation, and the precipitate was washed and suspended with 0.9% NaCl solution. 50% glutaraldehyde was added to an appropriate amount of suspension to reach a final concentration of 2.5%, fixing in a refrigerator at 4 °C for 4 h and then centrifuged (7500 rpm, 5 min), the final precipitate was washed three times with 0.9% NaCl solution. The suspension was then dehydrated using ethanol in a post-cellular gradient, and the suspension was dropped onto in silica foil and air-dried. For TEM, 1 mL samples were directly collected from the flasks, sealed, and stored in a 4°C refrigerator before analysis.

Valence analysis of the solid products was performed by X-ray photoelectron spectroscopy (XPS, ESCALAB250Xi, Thermo Fisher, UK). Sample was randomly selected from one of the triplicate flasks, centrifugating (10,000 rpm, 5 min), drying (80 °C, 24 h) and collected into a mortar for grinding. The same samples were also characterized by X-ray diffraction (XRD, Lab XRD-7000s, Shimadzu, Japan) with a scan speed of 6°/min and scanning range from 5°-90°.

## Results and discussion

### Te(IV) resistance and reducing characteristics of strain FF-1

Te(IV) was extremely toxic to most organisms, but it can be reduced by some microorganisms into Te^0^ or tellurides.^27^ Not surprisingly, the MIC for Te(IV) of strain FF-1 was as low as 10 μM, and the growth steady phase reached at about 60 h at this concentration (Fig.1a). The results of ICP showed a rapid decrease of the soluble Te concentration between 36 to 72 h, reaching the final removal percentage about 72% (Fig.1b). As the low Te(IV) concentration made the characterization of the solids products difficult, we tried to adding 1 mM Te(IV) after incubating strain FF-1 for 16 h. The high cell concentration (OD_660_ ~ 0.9) made the Te(IV) reduction at higher Te(IV) concentration became possible. After extra 30 h incubation, the solids was collected for XPS and XRD analysis. The results showed most of the Te species in solids were Te^0^ (accounting for 87.1%), and followed by Te(IV) (12.9%) (Fig. S1a). The XRD pattern also confirmed the existence of hexagonal structure of tellurium (PDF#36-1452) (Fig. S1b). Therefore, Te^0^ was confirmed as the main product of Te(IV) reduction by strain FF-1 as the other Te(IV) reducing bacteria did.^6, 28^ The residual Te(IV) in the solids may source from the Te(IV) adsorption at the surface of bacteria or BioTe^0^. On the other hand, no detectable methyl tellurides were found by GC after incubation for 168 h, and no homologous gene encoding for bacteria thiopurine methyltransferases was identified from the genome of strain FF-1. Therefore, the further Te^0^ reduction to the gaseous methyl tellurides species was unlikely to occur. To the best of our knowledge, no previous work mentioned one bacterium owing the Mn(II) oxidizing and Te(IV) reducing capacity simultaneously, although both functions has been widely reported in various bacterial lineage.^2, 14^ Therefore, *Bacillus* sp. FF-1 was an ideal candidate to investigate the possible reaction of *in-situ* produced BioMnOx and BioTe^0^.

**Figure 1.**
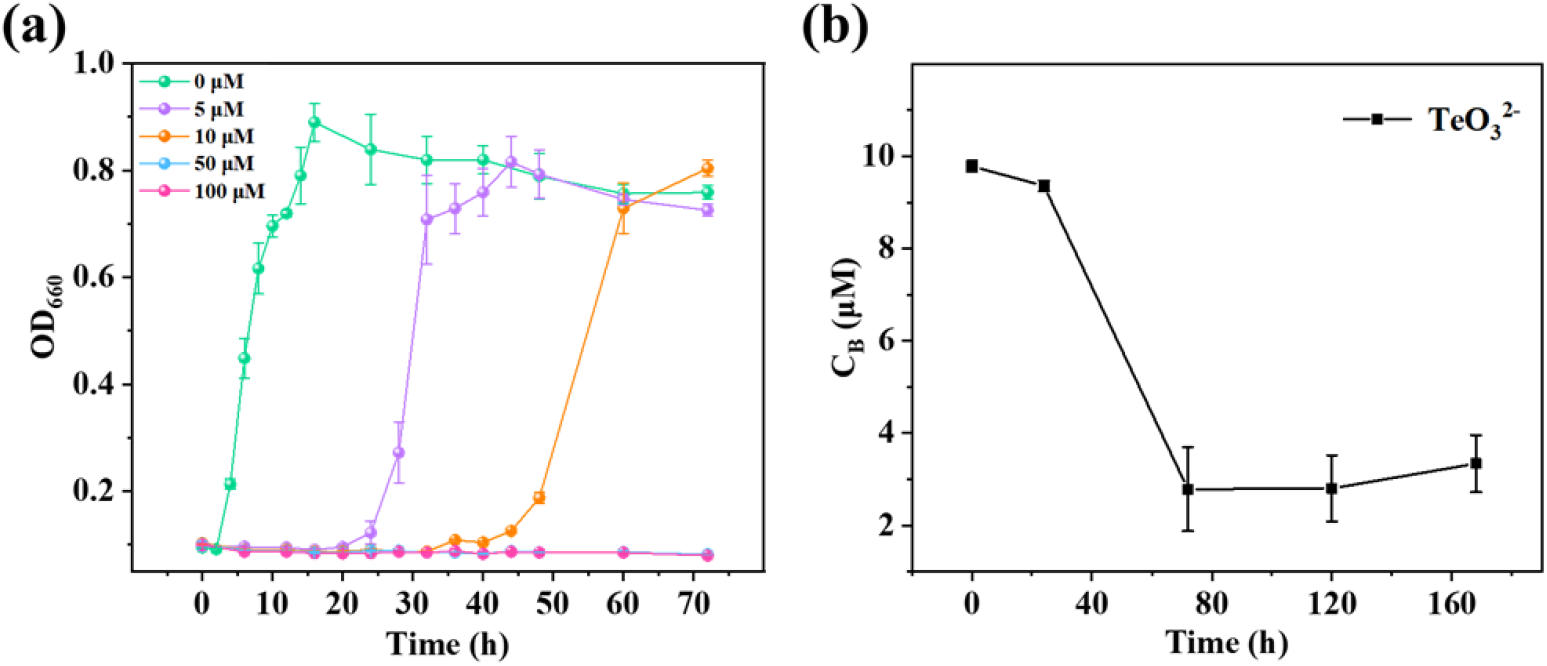
(a) Growth curve of strain FF-1 at different Te(IV) concentrations and (b) Te(IV) reduction curve of strain FF-1 with 10 μM Te(IV).

### Te conversion occurred in the presence of Mn(II) and Te(IV)

Three conditions were used to investigate the Te(IV) conversion by strain FF-1 in the presence of Mn(II) by changing the adding time of Mn(II) and Te(IV). Considering the trace amount of BioTe^0^ produced with 10 μM Te(IV), higher concentration of Mn(II) and Te(IV) was added in specific time intervals to obtained enough products for characterization. When 3 mM Mn(II) and 1 mM Te(IV) were added into the medium simultaneously (Mn0Te0h), a pale white precipitate appeared immediately and no color change was observed within 30 days. This phenomenon could also find in abiotic control without the bacterial cells, suggesting this reaction was a simple chemical process. It has been proved that Te(IV) could react with Fe(II) and Cr(III) to form Te(IV)-containing precipitates through the result of Te K-edge X-ray extended X-ray adsorption fine structure (EXAFs), thus Te(IV) cannot be reduced to Te^0^ in the presence of excess Fe(II) and Cr(III).^29, 30^ Therefore, the white precipitate may also the Mn-Te containing compounds, which may decrease the bioavailability of both ions. At the second condition, 3 mM Mn(II) was added at 0 h, while 1mM Te(IV) was added at 16 h (Mn0Te16h). In this condition, Mn(II) was firstly adsorbed on the bacterial cells and gradually be oxidized. After Te(IV) addition, the formation of white precipitates was also observed immediately, but the color of the medium became visually black after 24h incubation. The black color gradually lightens and finally became brown after 168 h incubation (Fig. S2a). This difference with the Mn0Te0h should be due to the pre-formation of BioMnOx before adding Te(IV). However, if 1 mM Te(IV) was added first after 16 h incubation of strain FF-1, and then Mn(II) added at 46 h (Te16Mn46h), no white precipitates was observed at all, and the solution color gradually changed from black to brown (Fig. S2b). The possible reason was most of the Te(IV) has been converted to Te^0^ or adsorbed on the surface of bacterial cells before adding Mn(II). The color change from black to brown in Mn0Te16h and Te16Mn46h suggested the possibility of Te^0^ (black) conversion by BioMnOx (brown) in these conditions.

To confirm the abovementioned hypothesis, the products at specific time intervals of these three conditions were characterized using XRD and XPS. Under the condition of Mn0Te16h, only significant Te(IV) peak was found over time in the XPS spectra of Te3d_5/2_, which was located at 575.3 eV (Fig. S3a).^31^ The XRD spectra for the samples collected at different time intervals also showed no evidence for the existence of Te^0^ (Fig. S3b). This result apparently controversial to the black color of medium appeared in this condition. The reason for that was most of the added Te(IV) may still co-precipitated with the residual Mn(II) because of the slow Mn(II) oxidation process (see below Fig. 3b). The BioTe^0^ produced from the free Te(IV) cannot be determined through the XPS spectra and XRD patterns due to the trace amount. For Mn0Te0h, the XPS of Mn3s and Te3d showed the dominant species were still Mn(II) and Te(IV), indicating no redox reaction occurred and the white precipitates were Mn(II)-Te(IV) containing compounds (Fig. S4a to S4c). The most reported Mn(II)-Te(IV) compound was MnTeO_3_, which usually synthesized by MnCl_2_ and Na_2_TeO_3_ and as theragnostic agents.^32^ However, there were no corresponding peaks were found in the XRD patterns of the Mn0Te0h before and after annealing at 200 °C, indicating a more complex products composition in our case (Fig. S4d). The co-removal of Cr(VI) and Te(IV) by *Shewanella oneidensis* MR-1 was also facilitated through co-precipitation, that Cr(VI) was reduced by bacteria, and fast complexed with Te(IV) to form Cr(III)-Te(IV) compounds.^29^ Under the condition of Te16Mn46h, XPS and XRD captured a more pronounced disappearance of the BioTe^0^. The peak area percentage of Te(0) decreased from 8.93% to 5.99% from 46 h to 214 h according to the XPS Te3d spectra (Fig. 2a). Meanwhile, the disappearance of Te(0) after adding Mn(II) could also confirmed by the XRD pattern (Fig. 2b). Therefore, the conversion of BioTe^0^ in Te16Mn46h could be confirmed.

**Figure 2.**
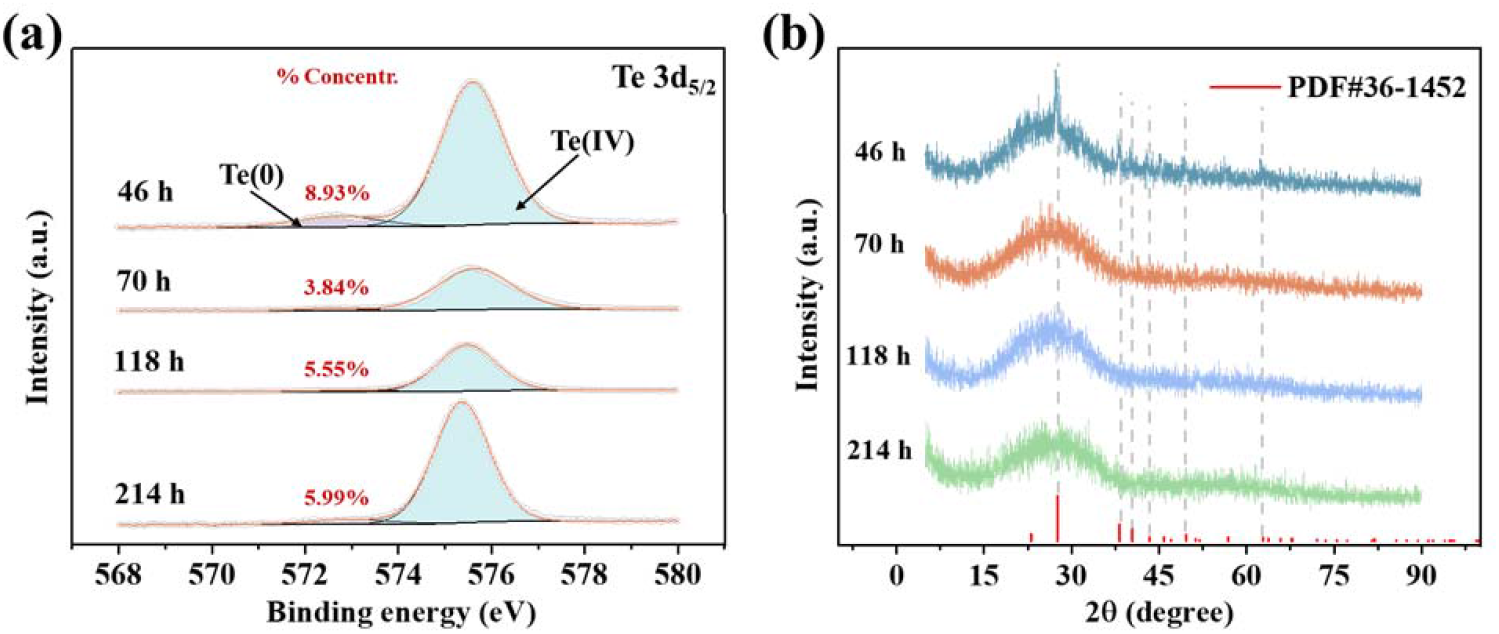
Characterization of XPS and XRD of samples during the reaction under “Te16Mn46h” conditions. The adding initial Te(IV) and Mn(II) concentrations were 1 mM and 5 mM, respectively.

**Figure 3.**
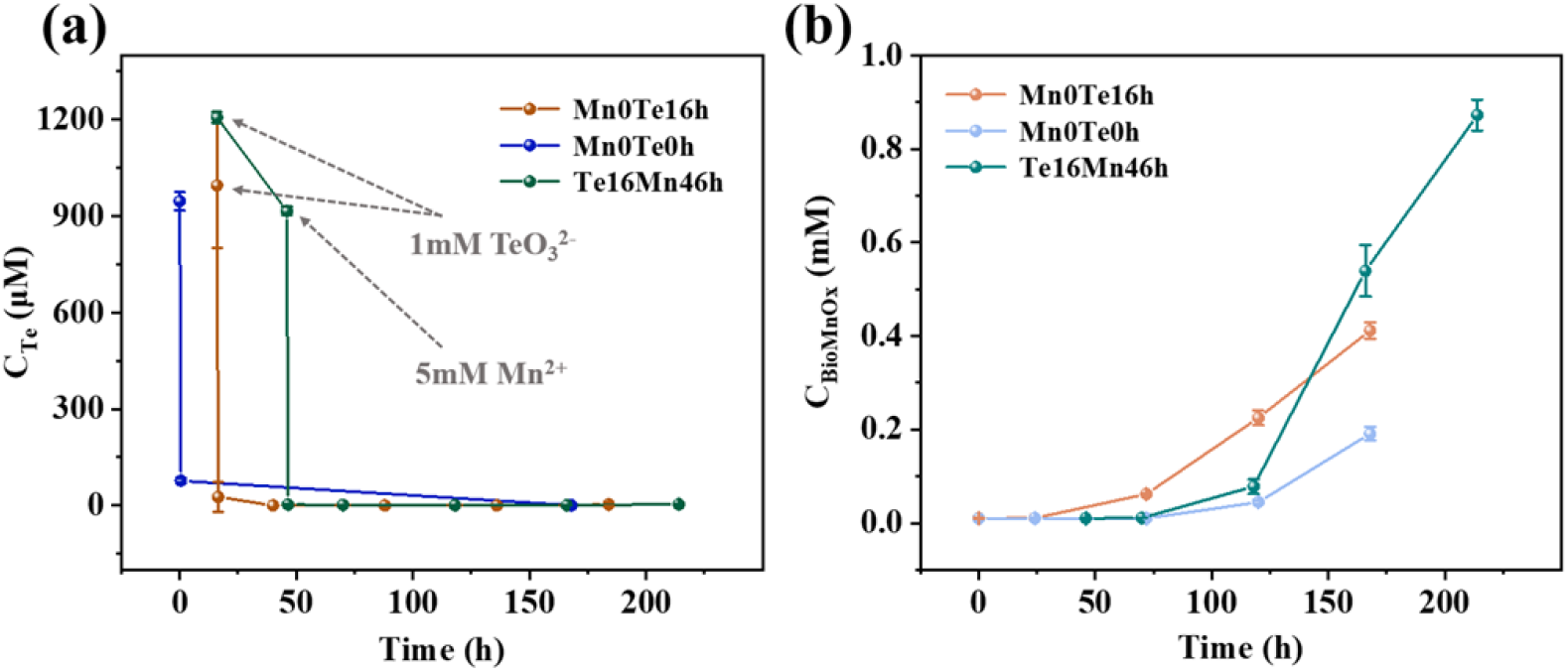
Variation curves of free Te concentration and BioMnOx content during the reaction under “Mn0Te16h; Mn0Te0h; Te16Mn46h” three conditions. The Mn(II) concentration in Mn0Te16h and Mn0Te0h were 3 mM, while in Te16Mn46h were 5 mM. All the Te(IV) concentration was 1 mM.

In addition, the changes of total Te concentration and manganese oxide content in the medium under the three conditions were measured. The total Te concentration were all decreased rapidly after adding Mn(II) into the medium, indicating the formation of Te-Mn compounds were the main fate of Te(IV) and Mn(II) in all three systems (Fig. 3a). The bioavailability of the Te(IV)-Mn(II) compounds was further evaluated by strain FF-1 and another manganese oxidizing bacterium HBX-1, that the manganese oxides produced by these two bacteria were all negligible even after 30 days (4.26 μM for strain HBX-1 and 2.64 μM for strain FF-1), indicating the low bioavailability of this compound. Therefore, the application of co-precipitation of Mn(II) and Te(IV) may help to facilitate quick and stable Te(IV) removal from wastewater than direct Te(IV) reduction to Te^0^. Also, it is reasonable to find the Mn(II) oxidation amounts of Mn0Te16h and Te16Mn46h were higher than Mn0Te0h due to the higher concentration of free Mn(II) (Fig. 3b). The continuous produced BioMnOx guarantee the enough oxidation power for BioTe^0^ reduction.

As enough BioTe^0^ formed in “Te16Mn46h” condition, The reaction between BioTe^0^ and BioMnOx could be further captured by TEM at different incubation time. Extracellular Te nanorods were firstly observed at 46 h before adding Mn(II), and the bacterial cells were coated with thick extracellular polymeric substances (Fig. S5a). This phenomenon suggested extracellular Te(IV) reduction was employed as the detoxification mechanism for strain FF-1 in the presence of high concentration of Te(IV). After the addition of Mn(II), many loose and porous structures were observed at the sample taken at 118 h incubation, which should be the Te(IV)-Mn(II) compound produced by the co-precipitation (Fig. S5b). At the same time, the rod-like structure gradually disassembly, and the size became smaller, verifying that the BioTe^0^ was unstable and underwent oxidation (Fig. S5c). The rod-like structure was barely observable in the sample after incubation for 214 h, instead by the small size sheet-like structure (Fig. S5d). Therefore, the conversion of Te(0) could also be supported by TEM. According to the previous reports, most Te(IV) reducing bacteria could form Te^0^ nanorods in the periplasmic and cytoplasmic space. However, adding lawsone or Fe(III) could promote the accumulation of extracellular Te^0^ in *Rhodobacter capsulatus* and *Shewanella oneidensi*s MR-1.^30, 33^ On the other hand, the bacterial formed manganese oxides generally coated at the surface of bacterial cells or released extracellularly.^34, 35^ The spatial interaction between extracellular BioTe^0^ and BioMnOx will be easier than the intracellular bioTe^0^, thus accelerate the possible Te^0^ oxidation by BioMnOx.

Considering the environmental-related concentration of Te(IV), the initial concentration of Te(IV) was further downregulated to 0.1 mM in “Te16Mn46h” condition. In this concentration, the BioTe^0^ oxidation became more obviously. 38.0% of the total Te(IV) were converted to BioTe^0^ when Mn(II) was added (46 h), according to the result of XPS. After 214 h incubation, the proportion of Te(0) decreased to 12.6% with the production of BioMnOx, indicating that the BioMnOx could oxidize the *in-situ* produced BioTe^0^ by an estimated rate of 0.151 μmol L^−1^ h^−1^ (Fig. 4). The bacterial tellurite reduction rates were much higher than this value, which could reach 20.45 μmol L^−1^ h^−1^ for *Raoultella* sp. WYA,^36^ 6 μmol L^−1^ h^−1^ for *Aromatoleum* sp. CIB,^37^ and 11.36 μmol L^−1^ h^−1^ for *Shinella* sp. WSJ-2.^38^ The nearly 100-folds lower rate of BioTe^0^ oxidation than BioTe^0^ reduction made the BioTe^0^ easily accumulate in most environment, but it may re-oxidized in the presence of Mn(II) and manganese oxidizing bacteria.

**Figure 4.**
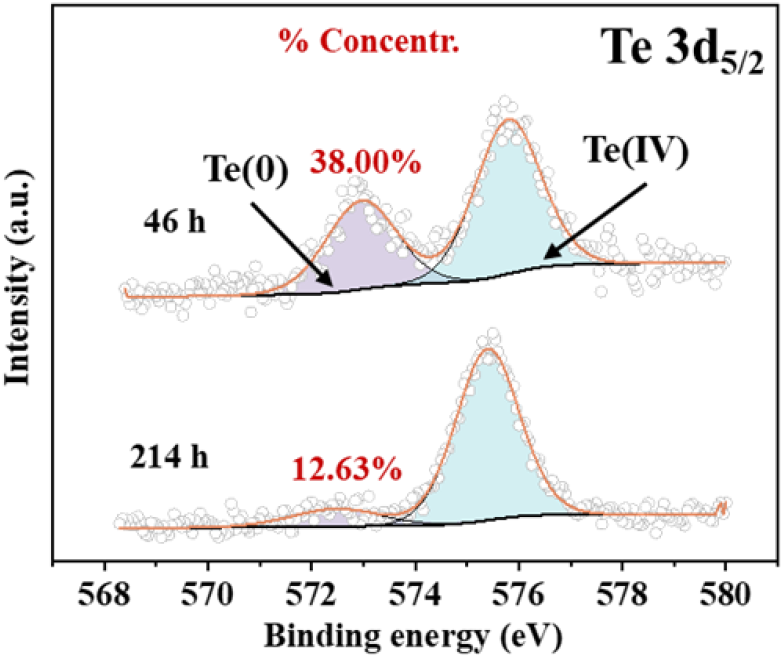
XPS characterization of the samples collected at the beginning (46 h) and the end of the reaction (214 h) under the condition of “Te16hMn46h” condition after adding Mn(II) (Te(IV)=0.1 mM, Mn(II)=5 mM)

### Te^0^ oxidation of chemical Te^0^ or BioTe^0^ produced by the other bacteria

As the biomolecules coated on the surface of biogenic nanoparticles may affect the oxidation of Te^0^, we examined the possibility of utilizing BioMnOx to oxidize chemical synthesized Te^0^ and BioTe^0^ produced by the other bacteria. When the chemical Te^0^ added into the BioMnOx-formed medium of strain FF-1, the distinct black color of Te^0^ gradually changed to brown during the reaction (Fig. 5a). The XPS spectrum showed the disappearance of Te(0) peak after 10 days’ incubation, while the proportion of the Te(0) in the initial sample was 34.46% (Fig. 5b). The calculated chemical Te^0^ oxidation by BioMnOx was 3.804 μmol L^−1^ d^−1^, which is much faster than the biomolecule-coated BioTe^0^. For the reaction of BioTe^0^ produced by another Te(IV)-reducing bacteria *Bacillus* sp. CC-1, the BioTe^0^ was added into the same BioMnOx-formed medium of strain FF-1. The XPS results showed that the signal peaks of Te(0) almost disappeared in all the conditions except sterilized BioTe^0^/sterilized BioMnOx (Fig. 5c). There was still 18.42% Te^0^ residual in the system of sterilized BioTe^0^/sterilized BioMnOx. Therefore, the BioTe^0^ oxidation by BioMnOx can occurred between different bacteria, and the continuously production of newly BioMnOx by manganese oxidizing bacteria may accelerate the BioTe^0^ oxidation.

**Figure 5.**
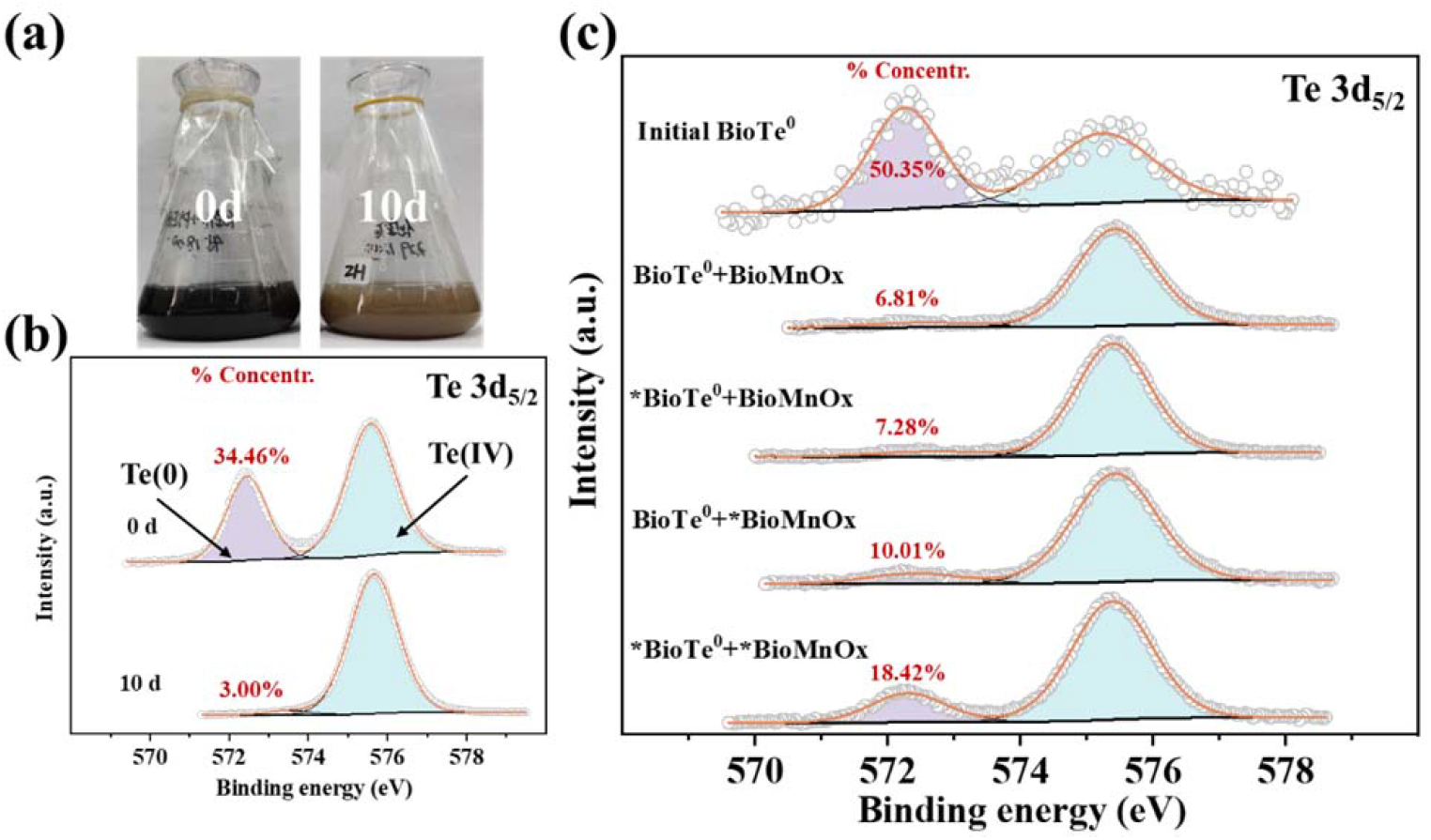
Color change (a) and XPS characterization (b) of the cultures containing chemogenic Te^0^ and BioMnOx-formed strain FF-1. (c) XPS spectra of the reaction products collected after reaction between BioTe^0^ from strain CC-1 and the BioMnOx produced from strain FF-1 for 14 days. (“*” represents that after high temperature sterilization)

The biogeochemical cycle of Te has been investigated for decades, and understand the environmental fate of metallic Te (Te^0^) is of great importance for Te(IV)-containing wastewater treatment since most of the final products was Te^0^. In previous study, the methylation process has been reported to consume Te^0^ and produced gaseous methyl tellurides species, thus questioned the stability of Te^0^ in natural environment.^39^ However, the Te^0^ methylation was much difficult than Te(IV) and Te(VI) methylation by thiopurine methyltransferases, and these enzymes were not widely existed in various bacteria. On the other hand, the strong acidic condition for possible Te^0^ oxidation by acidophilic bacterium were also uncommon in natural environment. Therefore, whether bacteria mediated Te^0^ oxidation could occur in neutral condition will great affect the fate of Te^0^.

In conclusion, in this study, we firstly found a bacteria possess both manganese-oxidizing and tellurite-reducing capacity, which was not reported before. According to the results, if Te(IV) and Mn(II) co-existed in the environment, this strain can convert them into Te(IV)-Mn(II) containing precipitates, which has high environmental stability and low bioavailability. Therefore, adding Mn(II) into Te(IV)-containing wastewater is an efficient approach to stabilize Te(IV) as solids. By this way, the Mn(II) oxidation and Te(IV) reduction rate will both become very slow in environment.

On the other hand, if BioTe^0^ has been formed, and then reacted with the *in-situ* formed BioMnOx by manganese oxidizing bacteria, it can be oxidized to Te(IV) again in a non-negligible rate. Therefore, the Te^0^ oxidation can truly occurred in neutral condition by the BioMnOx (Fig. 6a). Furthermore, the Te^0^ oxidation also occurred by the BioMnOx produced by strain FF-1 with chemical synthesized Te^0^, BioTe^0^ released into the environment, and the bacterial-bound BioTe^0^ (Fig. 6b-6d). These results further confirmed the possibility of BioMnOx mediated Te^0^ oxidation in a microecosystem, especially between different bacteria. Therefore, nevertheless the existence form of Te^0^, its stability can greatly affect by the existence of BioMnOx and MnOB. Therefore, the ecological risk of Te^0^ should be re-evaluated to design suitable Te(IV) removal processes.

**Figure 6.**
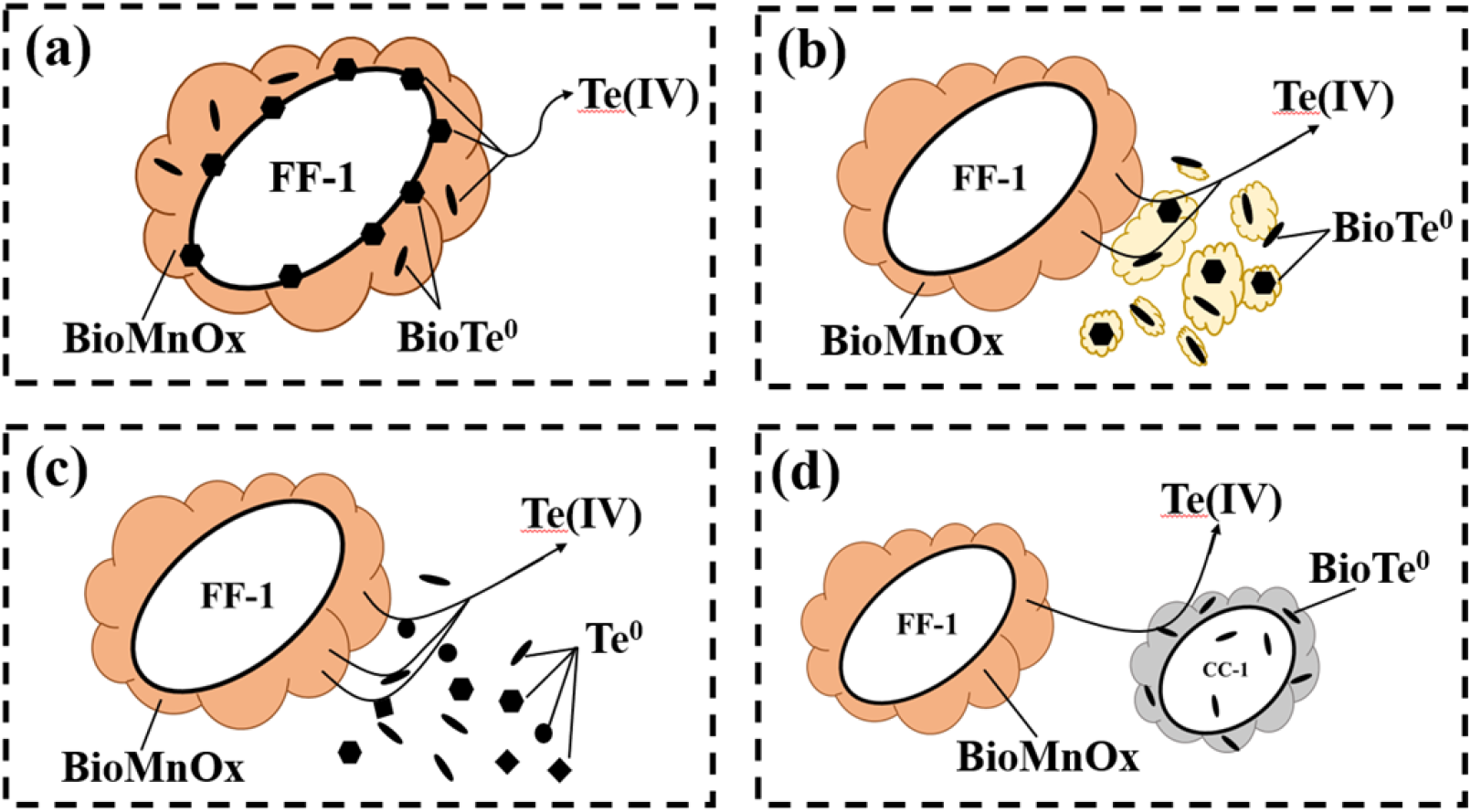
Mechanistic diagram of the reaction between BioMnOx of strain FF-1 and Te^0^ (a) BioMnOx and *in-situ* produced BioTe^0^ by strain FF-1; (b) BioMnOx and environmental BioTe^0^ (released from the lysis of other bacteria); (c) BioMnOx and chemical Te^0^; (d) BioMnOx and BioTe^0^-formed Te(IV)-reducing bacteria.

## Acknowledgements

The authors are financially supported by the National Natural Science Foundation of China (No. 41977197).

## References

Turner, R. J.; Borghese, R.; Zannoni, D., Microbial processing of tellurium as a tool in biotechnology. Biotechnol. Adv. 2012, 30 (5), 954–963.

Missen, O. P.; Ram, R.; Mills, S. J.; Etschmann, B.; Reith, F.; Shuster, J.; Smith, D. J.; Brugger, J., Love is in the Earth: A review of tellurium (bio)geochemistry in surface environments. Earth-Sci. Rev. 2020, 204, Article 103150.

Zheng, H. C.; Zhang, L.; Zhang, G. Y.; Gan, Y. H.; Xie, M.; Zhang, S. J., UV-Induced Redox Conversion of Tellurite by Biacetyl. Environ. Sci. Technol. 2021, 55 (24), 16646–16654.

Ramos-Ruiz, A.; Field, J. A.; Wilkening, J. V.; Sierra-Alvarez, R., Recovery of Elemental Tellurium Nanoparticles by the Reduction of Tellurium Oxyanions in a Methanogenic Microbial Consortium. Environ. Sci. Technol. 2016, 50 (3), 1492–1500.

Jin, W.; Su, J. L.; Chen, S. F.; Li, P.; Moats, M. S.; Maduraiveeran, G.; Lei, H., Efficient electrochemical recovery of fine tellurium powder from hydrochloric acid media via mass transfer enhancement. Sep. Purif. Technol. 2018, 203, 117–123.

Wang, Z. W.; Bu, Y. B.; Zhao, Y. H.; Zhang, Z. T.; Liu, L. F.; Zhou, H., Morphology-tunable tellurium nanomaterials produced by the tellurite-reducing bacterium *Lysinibacillus* sp. ZYM-1. Environ. Sci. Pollut. Res. 2018, 25 (21), 20756–20768.

Butz, Z. J.; Hendricks, A.; Borgognoni, K.; Ackerson, C. J., Identification of a TeO_3_^2-^ reductase/mycothione reductase from *Rhodococcus erythropolis* PR4. FEMS Microbiol. Ecol. 2021, 97 (1), Article 103150.

Presentato, A.; Turner, R. J.; Vasquez, C. C.; Yurkov, V.; Zannoni, D., Tellurite-dependent blackening of bacteria emerges from the dark ages. Environ. Chem. 2019, 16 (4), 266–288.

Mal, J.; Nancharaiah, Y. V.; Maheshwari, N.; van Hullebusch, E. D.; Lens, P. N. L., Continuous removal and recovery of tellurium in an upflow anaerobic granular sludge bed reactor. J. Hazard. Mater. 2017, 327, 79–88.

Shi, L. D.; Du, J. J.; Wang, L. B.; Han, Y. L.; Cao, K. F.; Lai, C. Y.; Zhao, H. P., Formation of nanoscale Te-0 and its effect on TeO_3_^2-^ reduction in CH_4_>-based membrane biofilm reactor. Sci. Total Environ. 2019, 655, 1232–1239.

Basnayake, R. S. T.; Bius, J. H.; Akpolat, O. M.; Chasteen, T. G., Production of dimethyl telluride and elemental tellurium by bacteria amended with tellurite or tellurate. Appl. Organomet. Chem. 2001, 15 (6), 499–510.

Prigent-Combaret, C.; Sanguin, H.; Champier, L.; Bertrand, C.; Monnez, C.; Colinon, C.; Blaha, D.; Ghigo, J. M.; Cournoyer, B., The bacterial thiopurine methyltransferase tellurite resistance process is highly dependent upon aggregation properties and oxidative stress response. Environ. Microbiol. 2012, 14 (10), 2645–2660.

Zhan, Y.; Shen, X.; Chen, M.; Yang, K.; Xie, H., Bioleaching of tellurium from mine tailings by indigenous *Acidithiobacillus ferrooxidans*. Lett. Appl. Microbiol. 2022, 75 (5), 1076–1083.

Zhou, H.; Fu, C., Manganese-oxidizing microbes and biogenic manganese oxides: characterization, Mn(II) oxidation mechanism and environmental relevance. Rev. Environ. Sci. Bio. 2020, 19 (3), 489–507.

Murray, K. J.; Tebo, B. M., Cr(III) is indirectly oxidized by the Mn(II)-Oxidizing bacterium *Bacillus* sp. strain SG-1. Environ. Sci. Technol. 2007, 41 (2), 528–533.

Murray, K. J.; Webb, S. M.; Bargar, J. R.; Tebo, B. M., Indirect oxidation of Co(II) in the presence of the marine Mn(II)-oxidizing bacterium *Bacillus* sp. strain SG-1. Appl. Environ. Microbiol. 2007, 73 (21), 6905–6909.

Ohnuki, T.; Ozaki, T.; Kozai, N.; Nankawa, T.; Sakamoto, F.; Sakai, T.; Suzuki, Y.; Francis, A. J., Concurrent transformation of Ce(III) and formation of biogenic manganese oxides. Chem. Geol. 2008, 253 (1-2), 23–29.

Yang, Y.; Ali, A.; Su, J.; Chang, Q.; Xu, L.; Su, L.; Qi, Z., Phenol and 17*β*-estradiol removal by *Zoogloea* sp. MFQ7 and in-situ generated biogenic manganese oxides: Performance, kinetics and mechanism. J. Hazard. Mater. 2022, 429, Article 128281.

Goff, J. L.; Wang, Y. W.; Boyanov, M. I.; Yu, Q.; Kemner, K. M.; Fein, J. B.; Yee, N., Tellurite Adsorption onto Bacterial Surfaces. Environ. Sci. Technol. 2021, 55 (15), 10378–10386.

Rosenfeld, C. E.; Sabuda, M. C.; Hinkle, M. A. G.; James, B. R.; Santelli, C. M., A Fungal-Mediated Cryptic Selenium Cycle Linked to Manganese Biogeochemistry. Environ. Sci. Technol. 2020, 54 (6), 3570–3580.

Che, L.; Xu, W. P.; Zhan, J. J.; Zhang, L.; Liu, L. F.; Zhou, H., Complete Genome Sequence of *Bacillus cereus* CC-1, A Novel Marine Selenate/Selenite Reducing Bacterium Producing Metallic Selenides Nanomaterials. Curr. Microbiol. 2019, 76 (1), 78–85.

Wu, J. H.; Kang, F.; Wang, Z. K.; Song, L.; Guan, X. Y.; Zhou, H., Manganese removal and product characteristics of a marine manganese-oxidizing bacterium *Bacillus* sp. FF-1. Int. Microbiol. 2022, 25 (4), 701–708.

Goff, J. L.; Boyanov, M. I.; Kemner, K. M.; Yee, N., The role of cysteine in tellurate reduction and toxicity. Biometals 2021, 34 (4), 937–946.

Tran, T. N.; Kim, D.-G.; Ko, S.-O., Synergistic effects of biogenic manganese oxide and Mn (II)-oxidizing bacterium *Pseudomonas putida* strain MnB1 on the degradation of 17 *α*-ethinylestradiol. J. Hazard. Mater. 2018, 344, 350–359.

Araya, M. A.; Swearingen, J. W.; Plishker, M. F.; Saavedra, C. P.; Chasteen, T. G.; Vásquez, C. C. J. J. J. o. B. I. C., *Geobacillus stearothermophilus* V *ubiE* gene product is involved in the evolution of dimethyl telluride in *Escherichia coli* K-12 cultures amended with potassium tellurate but not with potassium tellurite. J. Biol. Inorg. Chem. 2004, 9, 609–615.

Steinegger, M.; Soding, J., MMseqs2 enables sensitive protein sequence searching for the analysis of massive data sets. Nat. Biotechnol. 2017, 35 (11), 1026–1028.

Taylor, D. E., Bacterial tellurite resistance. Trends Microbiol. 1999, 7 (3), 111–115.

Cheng, M. M.; Sun, Y. Y.; Sui, X. R.; Zhang, H. K., Characterization of the differentiated reduction of selenite and tellurite by a halotolerant bacterium: Process and mechanism. J. Water Process. Eng. 2022, 47, Acticle 101016.

Kim, D. H.; Park, S.; Kim, M. G.; Hur, H. G., Accumulation of Amorphous Cr(III)-Te(IV) Nanoparticles on the Surface of *Shewanella oneidensis* MR-1 through Reduction of Cr(VI). Environ. Sci. Technol. 2014, 48 (24), 14599–14606.

Kim, D. H.; Kim, M. G.; Jiang, S. H.; Lee, J. H.; Hur, H. G., Promoted Reduction of Tellurite and Formation of Extracellular Tellurium Nanorods by Concerted Reaction between Iron and *Shewanella oneidensis* MR-1. Environ. Sci. Technol. 2013, 47 (15), 8709–8715.

Liu, M.; Xu, Y.; Liu, S. L.; Yin, S. L.; Liu, M. Y.; Wang, Z. Q.; Li, X. N.; Wang, L.; Wang, H. J., Phosphorus-modified ruthenium-tellurium dendritic nanotubes outperform platinum for alkaline hydrogen evolution. J. Mater. Chem. A 2021, 9 (8), 5026–5032.

He, Y.; Guo, Y. T.; Yu, Y. B.; Luo, P. F.; Qiu, H. L.; Huang, G. M., Facile Synthesis of Manganese Tellurite Nanoparticles for Magnetic Resonance Imaging-Guided Photothermal Therapy. Part. Part. Syst. Char. 2021, 38 (5), Article 101002.

Borghese, R.; Brucale, M.; Fortunato, G.; Lanzi, M.; Mezzi, A.; Valle, F.; Cavallini, M.; Zannoni, D., Extracellular production of tellurium nanoparticles by the photosynthetic bacterium *Rhodobacter capsulatus* (*Reprinted from Journal of Hazardous Materials*, vol 309, pg 202-209, 2016). J. Hazard. Mater. 2017, 324, 31–38.

Hosseinkhani, B.; Emtiazi, G., Synthesis and Characterization of a Novel Extracellular Biogenic Manganese Oxide (Bixbyite-like Mn_2_>O_3_>) Nanoparticle by Isolated *Acinetobacter* sp. Curr. Microbiol. 2011, 63 (3), 300–305.

Nelson, Y. M.; Lion, L. W.; Ghiorse, W. C.; Shuler, M. L., Production of biogenic Mn oxides by *Leprothrix discophora* SS-1 in a chemically defined growth medium and evaluation of their Pb adsorption characteristics. Appl. Environ. Microbiol. 1999, 65 (1), 175–180.

Nguyen, V. K.; Choi, W.; Ha, Y.; Gu, Y.; Lee, C.; Park, J.; Jang, G.; Shin, C.; Cho, S., Microbial tellurite reduction and production of elemental tellurium nanoparticles by novel bacteria isolated from wastewater. J. Ind. Eng. Chem. 2019, 78, 246–256.

Alonso-Fernandes, E.; Fernandez-Llamosas, H.; Cano, I.; Serrano-Pelejero, C.; Castro, L.; Diaz, E.; Carmona, M., Enhancing tellurite and selenite bioconversions by overexpressing a methyltransferase from *Aromatoleum* sp. CIB. Microb. Biotechnol., Article101111.

Wu, S. J.; Li, T. F.; Xia, X.; Zhou, Z. J.; Zheng, S. X.; Wang, G. J., Reduction of tellurite in *Shinella* sp. WSJ-2 and adsorption removal of multiple dyes and metals by biogenic tellurium nanorods. Int. Biodeterior. Biodegrad. 2019, 144, Article 101016.

Ollivier, P. R. L.; Bahrou, A. S.; Church, T. M.; Hanson, T. E., Aeration Controls the Reduction and Methylation of Tellurium by the Aerobic, Tellurite-Resistant Marine Yeast *Rhodotorula mucilaginosa*. Appl. Environ. Microbiol. 2011, 77 (13), 4610–4617.

